# Canonical neural networks perform active inference

**DOI:** 10.1101/2020.12.10.420547

**Authors:** Takuya Isomura, Hideaki Shimazaki, Karl Friston

## Abstract

This work considers a class of canonical neural networks comprising rate coding models, wherein neural activity and plasticity minimise a common cost function—and plasticity is modulated with a certain delay. We show that such neural networks implicitly perform active inference and learning to minimise the risk associated with future outcomes. Mathematical analyses demonstrate that this biological optimisation can be cast as maximisation of model evidence, or equivalently minimisation of variational free energy, under the well-known form of a partially observed Markov decision process model. This equivalence indicates that the delayed modulation of Hebbian plasticity—accompanied with adaptation of firing thresholds—is a sufficient neuronal substrate to attain Bayes optimal control. We corroborated this proposition using numerical analyses of maze tasks. This theory offers a universal characterisation of canonical neural networks in terms of Bayesian belief updating and provides insight into the neuronal mechanisms underlying planning and adaptive behavioural control.

## INTRODUCTION

The sentient behaviour of biological organisms is characterised by optimisation. Biological organisms recognise the state of their environment by optimising internal representations of the external (i.e., environmental) dynamics generating sensory inputs. In addition, they optimise their behaviour for adaptation to the environment, thereby increasing their probability of survival and reproduction. This biological self-organisation is typically formulated as the minimisation of cost functions [1–3], wherein a gradient descent on a cost function furnishes neural dynamics and synaptic plasticity. However, two fundamental issues remain to be established—namely, the characterisation of the dynamics of an arbitrary neural network as a generic optimisation process—and the correspondence between such neural dynamics and statistical inference [4] found in applied mathematics and machine learning. The present work addresses these issues by demonstrating that a class of canonical neural networks of rate coding models is functioning as— and thus universally characterised in terms of—variational Bayesian inference, under a particular but generic form of generative model.

Variational Bayesian inference offers a unified explanation for inference, learning, prediction, decision making, and the evolution of biological form [5,6]. This kind of inference rests upon a generative model that expresses a hypothesis about the generation of sensory inputs. Perception and behaviour can then be read as optimising the evidence for a ‘generative model’, inherent in sensory exchanges with the environment. The ensuing evidence lower bound (ELBO) [7], or equivalently variational free energy, then plays the role of a cost function. Variational free energy is the standard cost function in variational Bayes—and provides an upper bound on surprise (i.e., improbability) of sensory inputs. Minimisation of variational free energy, with respect to internal representations, then yields approximate posterior beliefs about external states. Similarly, the minimisation of variational free energy with respect to action on external states maximises the evidence or marginal likelihood of resulting sensory samples. This framework integrates perceptual (unsupervised), reward-based (reinforcement), and motor (supervised) learning in a unified formulation. In short, internal states of an autonomous system under a (possibly nonequilibrium) steady state can be viewed as parameterising posterior beliefs of external states [8–10]. In particular, active inference aims to optimise behaviours of a biological organism to minimise a certain kind of risk in the future [11–13], wherein risk is typically expressed in a form of expected free energy (i.e., the variational free energy expected under posterior predictive beliefs about the outcomes of a given course of action).

Crucially, as a corollary of the complete class theorem [14–16], any neural network minimising a cost function can be viewed as performing variational Bayesian inference, under some prior beliefs. We have previously introduced a reverse engineering approach that identifies a class of biologically plausible cost functions for neural networks [17]. We identified a class of cost functions for single-layer feedforward neural networks of rate coding models with a sigmoid (or logistic) activation function—based on the assumption that the dynamics of neurons and synapses follow a gradient descent on the same cost function. We subsequently demonstrated the mathematical equivalence between the class of cost functions for such neural networks and variational free energy under a particular form of generative model. This equivalence licenses variational Bayesian inference, and the ensuing free-energy principle, as the fundamental optimisation process underlying both dynamics and functions of such neural networks. Moreover, it enables one to characterise any variables and constants in the network in terms of quantities that play a role in variational Bayesian inference. However, it remains to be established whether active inference is an apt explanation for any given neural network interacting with the surrounding environment.

In most formulations, active inference goes further than simply assuming action and perception minimise variational free energy—it also considers the consequences of action as minimising expected free energy, i.e., planning as inference [18–22]. In what follows, we analytically and numerically demonstrate the implicit ability of neural networks to plan and minimise future risk, when viewed through the lens of active inference.

In the present work, we identify a class of biologically plausible cost functions for two-layer recurrent neural networks, under an assumption that neural activity and plasticity minimise a common cost function (referred to as assumption 1). Namely, we suppose a network of rate coding neurons with a sigmoid activation function, wherein the middle layer involves recurrent connections and the output layer provides feedback responses to the environment (assumption 2). In this work, such neural networks are referred to as canonical neural networks. Then, we demonstrate that the class of cost functions—describing their dynamics—can be cast as variational free energy under an implicit generative model, in the well-known form of a partially observable Markov decision process (POMDP). The gradient descent on the ensuing cost function naturally yields Hebbian plasticity [23–25] with an activity-dependent homeostatic term.

In particular, we consider the case where an arbitrary modulator [26–28] regulates synaptic plasticity with a certain delay (assumption 3) and demonstrate that such a modulation is identical to the update of a policy through a post-hoc evaluation of past decisions. The modulator renders the implicit cost function a risk function, which in turn renders behavioural control Bayes optimal—to minimise future risk. The proposed analysis affirms that active inference is an inherent property of canonical neural networks exhibiting delayed modulation of Hebbian plasticity. We discuss possible neuronal substrates that realise this modulation.

## RESULTS

### Overview of equivalence between neural networks and variational Bayes

First, we summarise the formal correspondence between neural networks and variational Bayes. A biological agent is formulated here as an autonomous system comprising a network of rate coding neurons. We presume that neural activity, action (decision), synaptic plasticity, and changes in any other free parameters minimise a common cost function *L* ≔ *L*(*o*_1:*t*_, *φ*) (c.f. assumption 1; **Fig. 1a**). Here, *o*_1:*t*_ ≔ {*o*_1_,…, *o*_*t*_} is a sequence of observations and *φ* ≔ {*x*_1:*t*_, *y*_1:*t*_, *W*, *ϕ*} is a set of internal states comprising the middle-layer (*x*_τ_) and output-layer (*y*_τ_) neural activity, synaptic strengths (*W*), and other free parameters (*ϕ*) that characterise *L* (e.g., firing threshold). Output-layer activity *y*_*t*_ determines the network’s actions or decisions *δ*_*t*_. Based on assumption 1, the update rule for the *i*-th component of *φ* is derived as the gradient descent on the cost function, 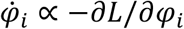. This determines the dynamics of neural networks, including their activity and plasticity.

**Figure 1.**
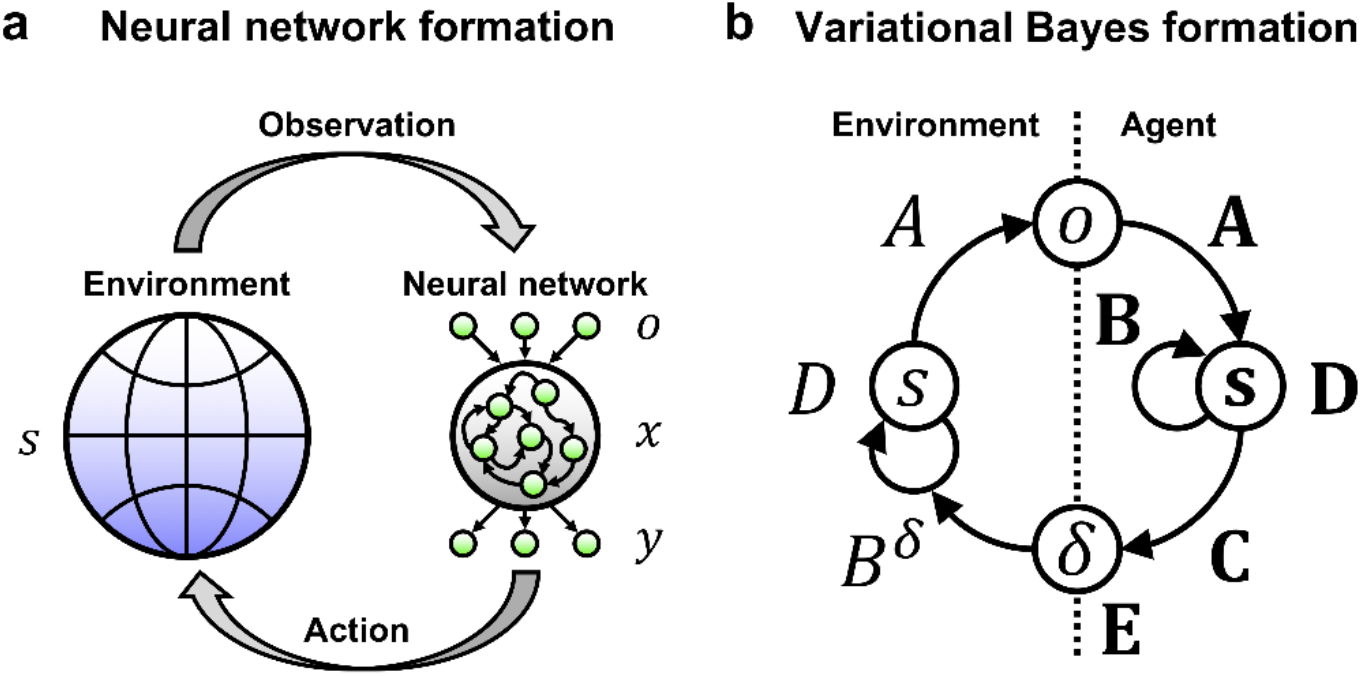
Schematic of an external milieu and neural network, and the corresponding Bayesian formation. (**a**) Interaction between the external milieu and autonomous system comprising a two-layer neural network. On receiving sensory inputs or observations *o*(*t*) that are generated from hidden states *s*(*t*), the network activity *x*(*t*) generates outputs *y*(*t*). The gradient descent on a neural network cost function *L* determines the dynamics of neural activity and plasticity. Thus, *L* is sufficient to characterise the neural network. The proposed theory affirms that the ensuing neural dynamics are self-organised to encode the posterior beliefs about hidden states and decisions. (**b**) Corresponding variational Bayesian formation. The interaction depicted in (a) is formulated in terms of a POMDP model, which is parameterised by *A*, *B*, *C* ∈ *θ* and *D*, *E* ∈ *λ*. Variational free energy minimisation allows an agent to self-organise to encode the hidden states of the external milieu—and to make decisions minimising future risk. Here, variational free energy *F* is sufficient to characterise the inferences and behaviours of the agent.

In contrast, variational Bayesian inference depicts a process of updating the prior distribution of external states *P*(*ϑ*) to the corresponding posterior distribution *Q*(*ϑ*) based on a sequence of observations. Here, *Q*(*ϑ*) approximates *P*(*ϑ*|*o*_1:*t*_). This process is formulated as a minimisation of the surprise of past-to-present observations—or equivalently maximisation of the model evidence—which is attained by minimising variational free energy as a tractable proxy. We suppose that the generative model *P*(*o*_1:*t*_, *ϑ*) is characterised by a set of external states, *ϑ* ≔ {*s*_1:*t*_, *δ*_1:*t*_, *θ*, *λ*}, comprising hidden states (*s*_τ_), decision (*δ*_τ_), model parameters (*θ*), and hyper parameters (*λ*) (**Fig. 1b**). Based on the given generative model, variational free energy is defined as a functional of *Q*(*ϑ*) as follows: *F*(*o*_1:*t*_, *Q*(*ϑ*)) ≔ E_*Q*(*ϑ*)_[− ln *P*(*o*_1:*t*_, *ϑ*) + ln *Q*(*ϑ*)]. Here, E_*Q*(*ϑ*)_[∙] ≔ ∫∙ *Q*(*ϑ*)*dϑ* denotes the expectation over *Q*(*ϑ*). In particular, we assume that *Q*(*ϑ*) is an exponential family and the posterior expectation of *ϑ*, **ϑ** ≔ E_*Q*(*ϑ*)_[*ϑ*], or its counterpart, are the sufficient statistics that parameterise (i.e., uniquely determine) *Q*(*ϑ*). Under this condition, *F* is reduced to a function of **ϑ**, *F* = *F*(*o*_1:*t*_, **ϑ**). The variational update rule for the *i*-th component of **ϑ** is given as the gradient descent on variational free energy, 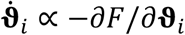.

Crucially, according to the complete class theorem, the dynamical system that minimises its cost function can be viewed as performing Bayesian inference under some generative model and prior beliefs. The complete class theorem [14–16] states that for any pair of admissible decision rules and cost functions, there is some generative model with prior beliefs that renders the decisions Bayes optimal. Thus, this theorem ensures the presence of a generative model that formally corresponds to the above-defined neural network characterised by *L*. Hence, this speaks to the equivalence between the class of neural network cost functions and variational free energy under such a generative model:

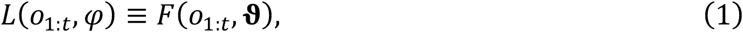

wherein the internal states of a network *φ* encode or parameterise the posterior expectation **ϑ**. This mathematical equivalence means that an arbitrary neural network, in the class under consideration, is implicitly performing active inference through variational free energy minimisation. Minimisation of *F* is achieved when and only when the posterior beliefs best match the true conditional probability of the external states. Thus, the dynamics that minimise *L* must induce a recapitulation of the external states in the internal states of the neural network. This is a fundamental aspect of optimisation in neural networks. This notion is essential to understand the functional meaning of the dynamics evinced by an arbitrary neural network, which is otherwise unclear by simply observing the network dynamics.

Note that being able to characterise the neural network in terms of maximising model evidence lends it an ‘explainability’, in the sense that the internal (neural network) states and parameters encode Bayesian beliefs or expectations about the causes of observations. In other words, the generative model explains how outcomes were generated. However, the complete class theorem does not specify the form of generative model for any given neural network. To address this issue, we formulate active inference using a particular form of POMDP models, whose states take binary values. This facilitates identification of a class of generative models that corresponds to a class of canonical neural networks—comprising rate coding models with the sigmoid activation function.

### Active inference formulated using a postdiction of past decisions

In this section, we define generative model and ensuing variational free energy that correspond to a class of canonical neural networks that will be considered in the subsequent section. The external milieu is expressed as a discrete state space in the form of a POMDP (**Fig. 2**). The generation of observations 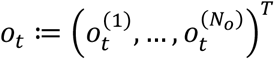 from external or hidden states milieu 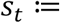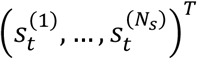 is expressed in the form of a categorical distribution, *P*(*o*_*t*_ |*S*_*t*_, *A*) = Cat(*A*), where *A* is referred to as the likelihood mapping. Our agent receives *o*_*t*_, infers latent variables (hidden states) *S*_*t*_, and provides a feedback decision 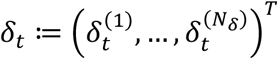 to the external milieu. Thus, the state transition at time *t* depends on *δ*_*t*-1_, characterised by the state transition matrix *B*^δ^, *P*(*S*_*t*_|*S*_*t*-1_, *δ*_*t*-1_, *B*) = Cat(*B*^δ^). Each element of *S*_*t*_, *o*_*t*_, and *δ*_*t*_ adopts a binary value, which is suitable for characterising generative models implicit in canonical neural networks (see below). When dealing with external states that factorize (e.g., *what* and *where*), block matrices *A* and *B* (and *C*) are the outer products of sub matrices (please refer to Methods A for further details); see also [17]. Hence, we define the generative model as follows:

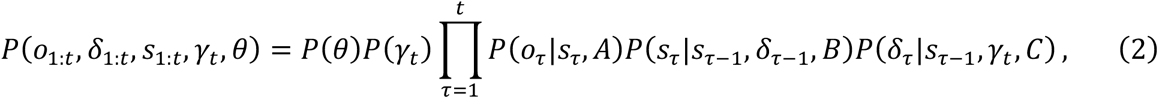

where 1 ≤ *τ* ≤ *t* denotes an arbitrary time in the past, *θ* ≔ {*A*, *B*, *C*} constitute the set of parameters, and *γ*_*t*_ is the current risk (see below). Note that initial states and decisions are characterised by prior distributions *P*(*S*_1_) = Cat(*D*) and *P*(*δ*_1_) = Cat(*E*), where *D* and *E* are block vectors.

**Figure 2.**
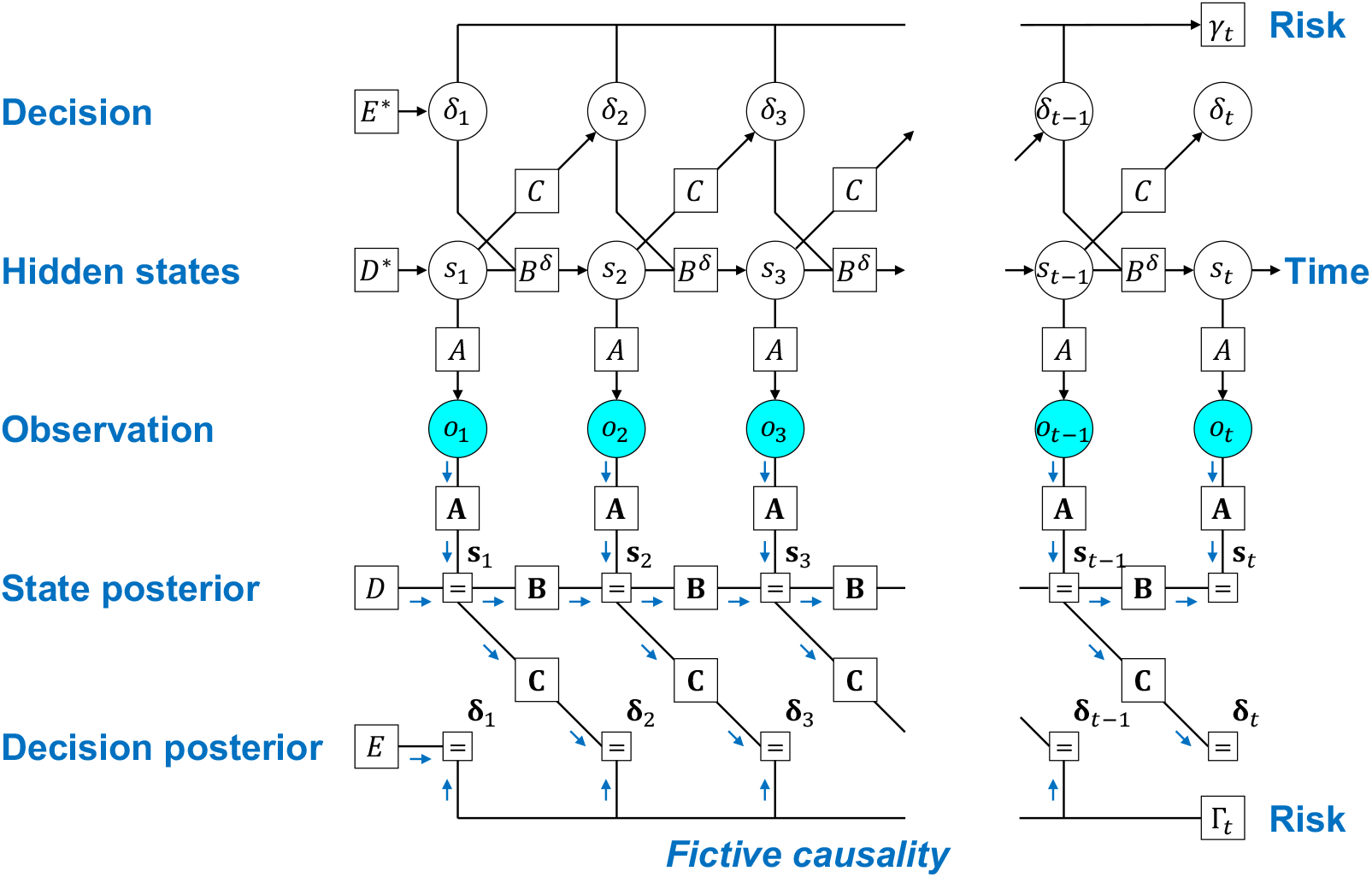
Factor graph depicting a fictive causality of factors that the generative model hypothesises. The POMDP model is expressed as a Forney factor graph [48,49] based upon the formulation in [50]. The arrow from the present risk *γ*_*t*_—sampled from Γ_*t*_—to a past decision *δ*_τ_ optimises the policy in a post-hoc manner, to minimise future risk. In reality, the current error *γ*_*t*_ is determined based on past decisions (top). In contrast, decision making to minimise the future risk implies a fictive causality from *γ*_*t*_ to *δ*_*t*_ (bottom). Inference and learning correspond to the inversion of this generative model. Postdiction of past decisions is formulated as the learning of the policy mapping, conditioned by *γ*_*t*_. Here, *A*, *B*, and *C* indicate matrices of the conditional probability, and bold case variables are the corresponding posterior beliefs. Moreover, *D*\* and *E*\* indicate the true prior beliefs about hidden states and decisions, while *D* and *E* indicate the priors that the network operates under. When and only when *D* = *D*\* and *E* = *E*\*, inferences and behaviours are optimal for a given task or set of environmental contingencies, and biased otherwise.

The agent makes decisions to minimise a risk function Γ_*t*_ ≔ Γ(*o*_1:*t*_, **s**_1:*t*_, **δ**_1:*t*-1_, **θ**) that it employs (where 0 ≤ Γ_*t*_ ≤ 1). Because the current risk Γ_*t*_ is a consequence of past decisions, the agent needs to select decisions that minimise the future risk. In this sense, Γ_*t*_ corresponds to the expected free energy in the usual formulation of active inference [12,13].

In our POMDP model, we formulate the operation to optimise decisions using a fictive causality from the current risk Γ_*t*_ to past decisions *δ*_1_,…, *δ*_*t*-1_ (**Fig. 2**). Although this is not the true causality in the real generative process that generates sensory data, we intend to model the manner that an agent subjectively evaluates its previous decisions after experiencing their consequences. This fictive causality is expressed in the form of a categorical distribution,

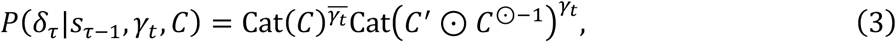

wherein policy mapping *C* is switched by a binarized risk *γ*_*t*_ ∈ {0,1}—sampled from 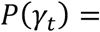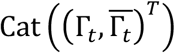—in a form of mixture model^*^. Note that *C*′ denotes a normalisation factor that is negligible in the following formulations, ⊙ indicates the element-wise product operator, and *C*^⊙-1^ means the element-wise inverse of matrix *C*. Throughout the manuscript, the overline variable indicates one minus the variable (e.g., 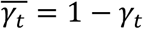). This model presumes that a past decision *δ*_τ_ (1 ≤ *τ* ≤ *t* − 1) is determined based on a past state *S*_τ-1_ and the current risk *γ*_*t*_. In contrast, the current decision *δ*_*t*_ is determined to minimise the future risk, *P*(*δ*_*t*_|*S*_*t*-1_, *C*) = Cat(*C*), because the agent has not yet observed the consequences of the current decision.

Importantly, the agent needs to keep selecting ‘good’ decisions while avoiding ‘bad’ decisions. To this end, we suppose that the agent learns from the failure of decisions, by assuming that the bad decisions were sampled from the opposite of the optimal policy mapping. In other words, the agent is assumed to have the prior belief such that the decision—sampled from Cat(*C*)—should result in *γ*_*t*_ = 0, while sampling from Cat(*C*′ ⊙ *C*^⊙-1^) should yield *γ*_*t*_ = 1. This construction enables the agent to conduct a postdiction of its past decisions—and thereby to update the policy mapping to minimise future risk—by associating the past decision rule (policy) with the current risk. In the next section, we will explain the biological plausibility of this form of adaptive behavioural control, wherein the update of the policy mapping turns out to be identical to a delayed modulation of Hebbian plasticity.

Variational Bayesian inference refers to the process that optimises the posterior belief *Q*(*ϑ*). Based on the mean-field approximation, *Q*(*ϑ*) is expressed as

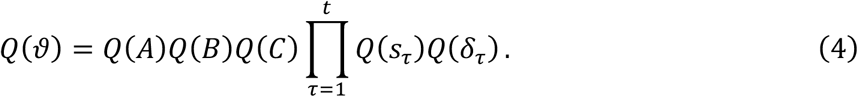

Here, the posterior beliefs about states and decisions are categorical probability distributions, *Q*(*S*_τ_) = Cat(**s**_τ_) and *Q*(*δ*_τ_) = Cat(**δ**_τ_), whereas those about parameters are Dirichlet distributions, *Q*(*A*) = Dir(**a**), *Q*(*B*) = Dir(**b**), and *Q*(*C*) = Dir(**c**). Throughout the manuscript, bold case variables (e.g., **s**_τ_) denote the posterior expectations of the corresponding italic case random variables (e.g., *S*_τ_). The agent samples a decision *δ*_τ_ at time *t* from the posterior distribution *Q*(*δ*_τ_). In this paper, the posterior belief of transition mapping is averaged over all possible decisions, **B** = E_Q(δ)_[**B**^δ^], to ensure the exact correspondence to canonical neural networks (see below). We use **θ** ≔ {**a**, **b**, **c**} to denote the parameter posteriors. For simplicity, here we suppose that state and decision priors (*D*, *E*) are fixed.

Under the above-defined generative model and posterior beliefs, the ensuing variational free energy is analytically expressed as follows:

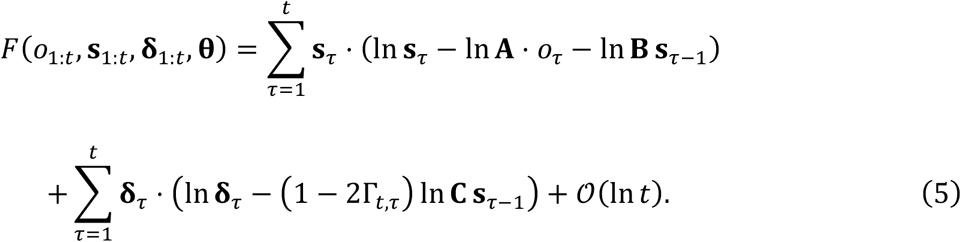

The derivation details are provided in Methods B. Note that Γ_*,τ*_ = 0 for *τ* = *t*; otherwise, Γ_*,τ*_ = Γ_*t*_. The order ln *t* term indicates the complexity of parameters, which is negligible when the leading order term is large. The gradient descent on variational free energy updates the posterior beliefs about hidden states (**s**_*t*_), decisions (**δ**_*t*_), and parameters (**θ**). The optimal posterior beliefs that minimise variational free energy are obtained as the fixed point of the implicit gradient descent, which ensures that ∂*F*/∂**s**_*t*_ = 0, ∂*F*/∂**δ**_*t*_ = 0, and ∂*F*/∂**θ** = *O*. The explicit forms of the posterior beliefs are provided in Methods C.

To explicitly demonstrate the formal correspondence with the cost functions for neural networks considered below, we further transform the variational free energy: based on Bayes theorem *P*(*S*_τ_|*S*_τ-1_, *B*^δ^) ∝ *P*(*S*_τ-1_|*S*_τ_, *B*^δ^)*P*(*S*_τ_), the inverse transition mapping is expressed as **B**^✝^ = **B**^*T*^diag[*D*]^-1^ using the state prior *P*(*S*_τ_) = Cat(*D*) (where *P*(*S*_τ-1_) is supposed to be a flat prior belief). Moreover, from *P*(*δ*τ|*S*_τ-1_, *γ*_*t*_, *C*) ∝ *P*(*S*_τ-1_|*δ*_τ_, *γ*_*t*_, *C*)*P*(*δ*_τ_), the inverse policy mapping is expressed as **C**^✝^ = **C**^*T*^ diag[*E*]^-1^ using the decision prior *P*(*δ*_*t*_) = Cat(*E*). Using these relationships, equation (5) is transformed into the form shown in **Fig. 3**(top). Please see Methods B for further details. This specific form of variational free energy constitutes a class of cost functions for canonical neural networks, as we will see below.

**Figure 3.**
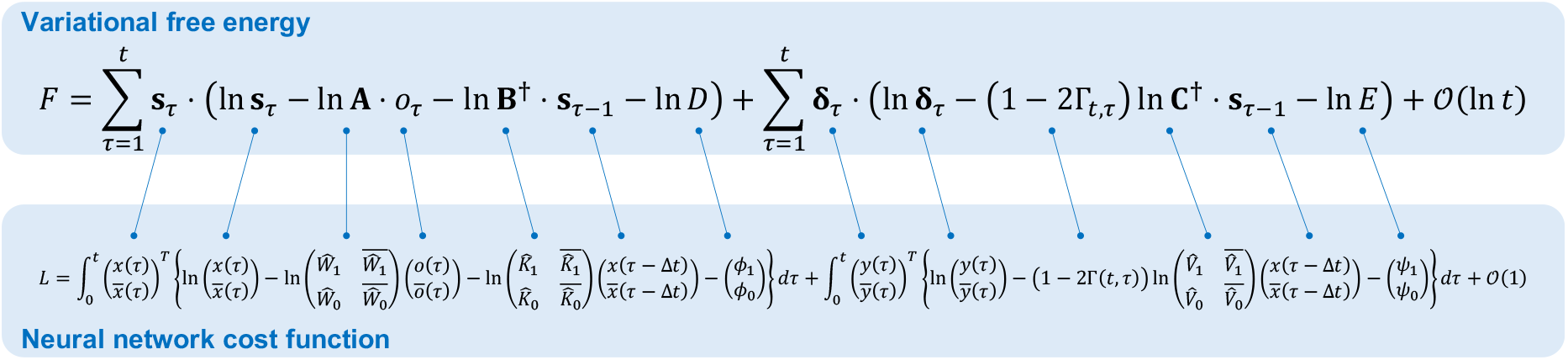
Mathematical equivalence between variational free energy and neural network cost functions, depicted by one-by-one correspondence of their components. Top: variational free energy transformed from equation (5) using the Bayes theorem. Here, **B**^✝^ = **B**^_^diag[*D*]^*t*-1^ and **C**^✝^ = **C**^_^diag[*E*]^*t*-1^ indicate the inverse mappings, and *D* and *E* are the state and decision priors. Bottom: neural network cost function that is a counterpart to the aforementioned variational free energy. In this equation, 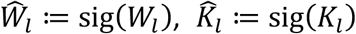, and 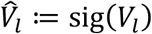 (for *l* = 0,1) indicate the sigmoid functions of synaptic strengths. Moreover, *ϕ*_*l*_ and φ_*l*_ are perturbation terms that characterise the bias in firing thresholds. Here, 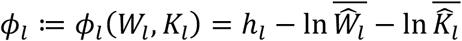 is a function of *W*_*l*_ and *K*_*l*_, while 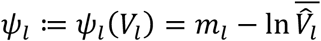 is a function of *V*_*l*_. When 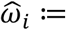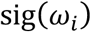 is the sigmoid function of ω_*i*_, 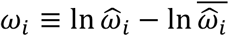 holds for an arbitrary ω_*i*_. Using this relationship, equation (7) is transformed into the form presented at the bottom of this figure. This form of cost functions formally corresponds to variational free energy expressed on the top of this figure.

In summary, variational free energy minimisation underwrites optimisation of posterior beliefs. In neurobiological formulations, it is usually assumed that neurons encode **s**_*t*_ and **δ**_*t*_, while synaptic strengths encode **θ** [12,13]. In what follows, we demonstrate that the internal states of canonical neural networks encode posterior beliefs.

### Canonical neural networks perform active inference

In this section, we identify the neuronal substrates that correspond to components of the active inference scheme defined above. We consider a class of two-layer neural networks with recurrent connections in the middle layer (**Fig. 1a**). The modelling of the networks in this section (referred to as canonical neural networks) is based on the following three assumptions—that reflect physiological knowledge: (1) a gradient descent on a cost function *L* determines the updates of neural activity and synaptic weights; (2) neural activity is updated by the weighted sum of inputs, and its fixed point is expressed in a form of the sigmoid (or logistic) function; and (3) a modulatory factor mediates synaptic plasticity in a post-hoc manner.

Based on assumption 2, we formulate neural activity in the middle layer (*x*) and output layer (*y*) as follows:

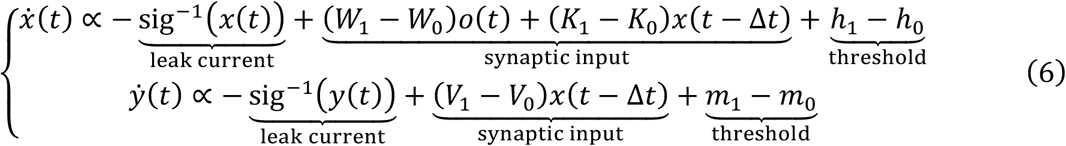

Here, 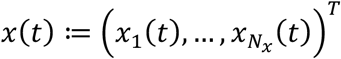 and 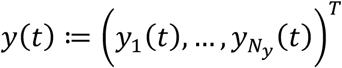 denote column vectors of firing intensities; 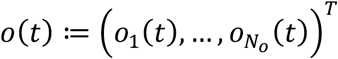 is a column vector of binary sensory inputs; *W*_1_, 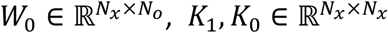, and *V*_1_, 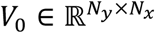 are synaptic strength matrices; and *h*_1_ ≔(*W*_1_, *K*_1_), *h*_0_ ≔ (*W*_0_, *K*_0_), *m*_1_ ≔ *m*_1_(*V*_1_), and *m*_0_ ≔ *m*_0_(*V*_0_) are adaptive firing thresholds that depend on synaptic strengths.

One may think of *W*_1_, *K*_1_, and *V*_1_ as excitatory synapses, whereas *W*_0_, *K*_0_, and *V*_0_ can be regarded as inhibitory synapses. Here, (*W*_1_ − *W*_0_)*O*(*t*) represents the total synaptic input from the sensory layer, and (*K*_1_ − *K*_0_)*x*(*t* − Δ*t*) forms a recurrent circuit with a time delay Δ*t* > 0. Receiving inputs from the middle layer *x*(*t*), the output-layer neural activity *y*(*t*) determines the decision 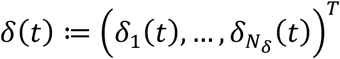, i.e., Prob[*δ*_*i*_(*t*) = 1] = *y*_*i*_(*t*). We select the inverse sigmoid (i.e., logit) leak current to ensure that the fixed point of equation (6) (i.e., *x* and *y* that ensure 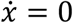 and 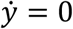) has the form of a sigmoid activation function (c.f. assumption 2). The sigmoid activation function is also known as the neurometric function [29].

Without loss of generality, equation (6) can be cast as the gradient descent on cost function *L*. Such a cost function can be identified by simply integrating the right-hand side of equation (6) with respect to *x* and *y*, consistent with previous treatments [17]. Moreover, we presume that output-layer synapses (*V*_1_ and *V*_0_) are updated through synaptic plasticity mediated by the modulator Γ(*t*) (c.f. assumption 3; 0 ≤ Γ(*t*) ≤ 1), as a model of plasticity modulations that are empirically observed [26–28]. Because neural activity and synaptic plasticity minimise the same cost function *L*, the derivatives of *L* must generate the modulated synaptic plasticity. Under these constraints reflecting assumptions 1–3, a class of cost functions is identified as follows:

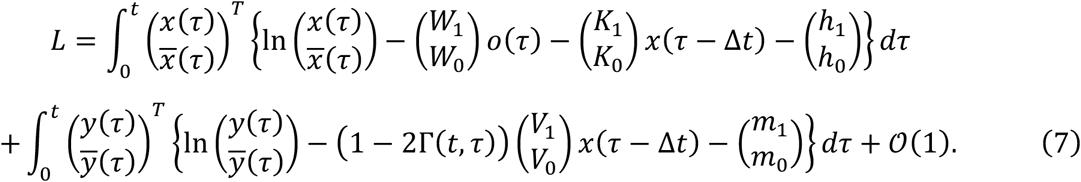

Here, 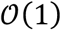—that denotes a function of synaptic strengths—is of a smaller order than the other terms that are of order *t*. Thus, 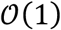 is negligible when *t* is large. We suppose Γ(*t*, *τ*) = 0 for *t* − Δ*t* < *τ* ≤ *t* and Γ(*t*, *τ*) = Γ(*t*) for 0 ≤ *τ* ≤ *t* − Δ*t*, to satisfy assumptions 1–3. This means that the optimisation of *L* by associative plasticity is mediated by Γ(*t*). Note that a gradient descent on *L*, i.e. 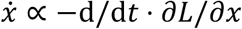 and 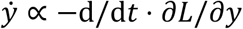, has the same functional form (and solution) as equation (6).

Synaptic plasticity rules conjugate to the above rate coding model can now be expressed as a gradient descent on the same cost function *L*, according to assumption 1. To simplify notation, we define synaptic strength matrix as ω_*i*_ ∈ {*W*_1_, *W*_0_, *K*_1_, *K*_0_, *V*_1_, *V*_0_}, pre-synaptic activity as *pre*_*i*_(*t*) ∈ {*o*(*t*), *o*(*t*), *x*(*t* − Δ*t*), *x*(*t* − Δ*t*), *x*(*t* − Δ*t*), *x*(*t* − Δ*t*)}, post-synaptic activity as 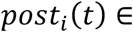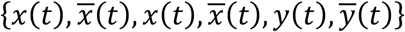, and firing thresholds as *n*_*i*_ ∈ {*h*_1_, *h*_0_, *h*_1_, *h*_0_, *m*_1_, *m*_0_}. Thus, synaptic plasticity in the first layer (*i* = 1,…,4) is derived as follows:

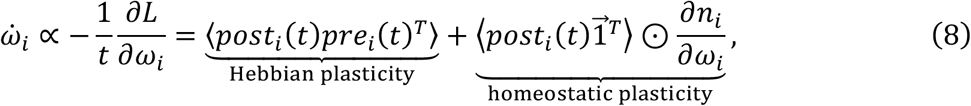

Moreover, synaptic plasticity in the second layer (*i* = 5,6) is derived as follows:

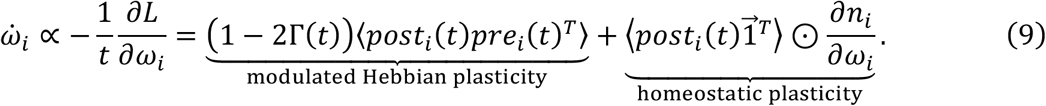

Here, 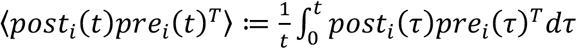 indicates the average over time and 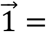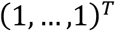 is a vector of ones.

These synaptic update rules are biologically plausible as they comprise Hebbian plasticity— determined by the outer product of pre- and post-synaptic activity—accompanied by an activity-dependent homeostatic term. In equation (9), the neuromodulator Γ(*t*)—that encodes an arbitrary risk—alters the form of Hebbian plasticity in a post-hoc manner. This can facilitate the association between past decisions and the current risk, thus leading to the optimisation of the decision rule to minimise future risk. In short, Γ(*t*) > 0.5 yields Hebbian plasticity, whereas Γ(*t*) < 0.5 yields anti-Hebbian plasticity. Empirical observations suggest that some modulators [26–28], such as dopamine neurons [30], are a possible neuronal substrate of Γ(*t*); please see

### Discussion for further details

Based on the above considerations, we now establish the formal correspondence between the neural network cost function and variational free energy. Under the aforementioned three minimal assumptions, we identify the neural network cost function as equation (7). Equation (7) can be transformed into the form shown in **Fig. 3**(bottom) using sigmoid functions of synaptic strengths (e.g., 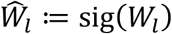 for *l* = 0,1). Here, the firing thresholds (*h*_*l*_, *m*_*l*_) are replaced with the perturbation terms in the thresholds, 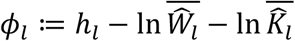 and 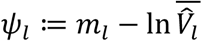. **Figure 3** depicts the formal equivalence between the neural network cost function (**Fig. 3**, bottom) and variational free energy (**Fig. 3**, top), visualised by one-by-one correspondence between their components. The components of variational free energy—including the log likelihood function and complexities of states and decisions—re-emerge in the neural network cost function.

This means that when 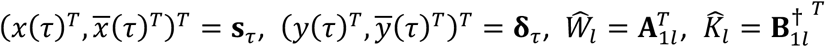, and 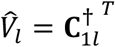 (for *l* = 0,1) the neural network cost function is identical to variational free energy, up to the negligible ln *t* residual. This further endorses the asymptotic equivalence of equations (5) and (7).

The neural network cost function is characterised by the perturbation terms implicit in firing thresholds *ϕ* ≔ (*ϕ*_1_, *ϕ*_0_) and φ ≔ (φ_1_, φ_0_). These terms correspond to the state and decision priors, ln *P*(*S*_*t*_) = ln *D* = *ϕ* and ln *P*(*δ*_*t*_) = ln *E* = φ, respectively. Hence, this class of cost functions for canonical neural networks is formally homologous to variational free energy, under the particular form of POMDP generative model, defined in the previous section. In other words, equations (2) and (5) express the class of generative models—and ensuing variational free energy—that ensure equation (1) is apt, for the class of canonical neural networks considered. This in turn suggests that any canonical neural network in this class is implicitly performing active inference. **Table 1** summarises the correspondence between the quantities of the neural network and their homologues in variational Bayes.

**Table 1.**
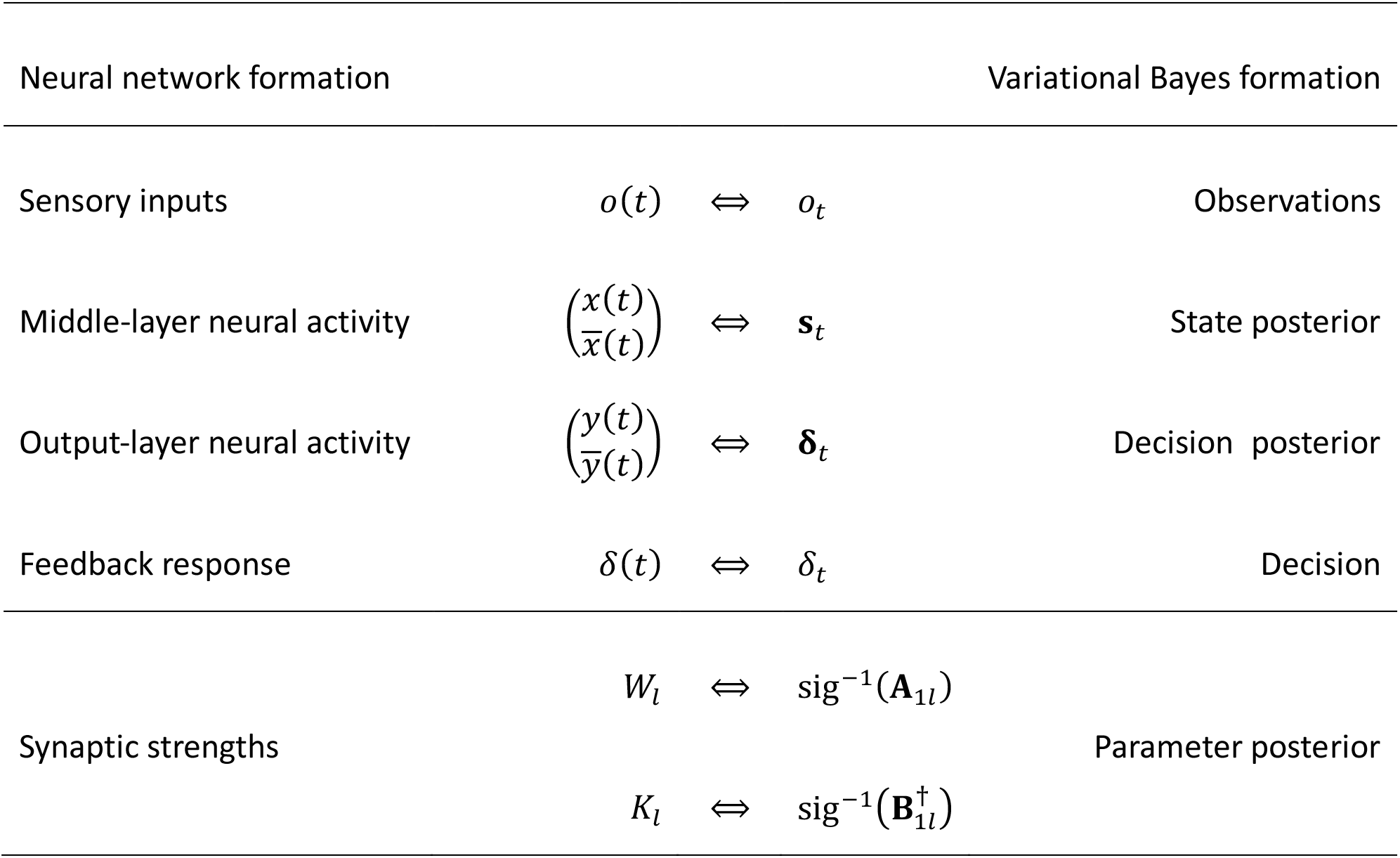
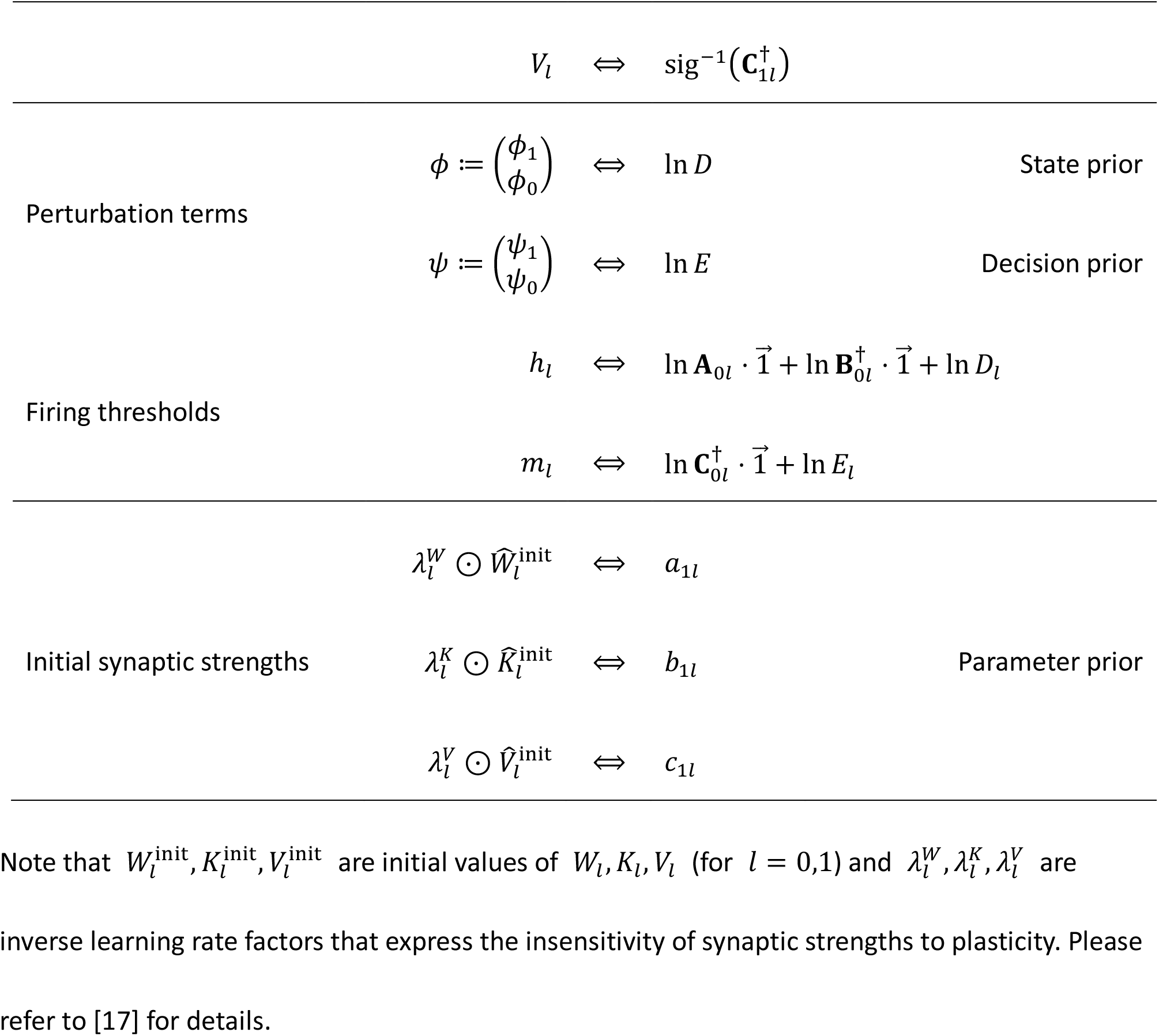
Correspondence of variables and functions.

In summary, when a neural network minimises the cost function with respect to its activity and plasticity, the network self-organises to furnish responses that minimise a risk implicit in the cost function. This biological optimisation is identical to variational free energy minimisation under a particular form of POMDP model. Hence, this equivalence indicates that minimising the expected risk through variational free energy minimisation is an inherent property of canonical neural networks featuring a delayed modulation of Hebbian plasticity.

Note that 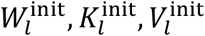 are initial values of *W*_*l*_, *K*_*l*_, *V*_*l*_ (for *l* = 0,1) and 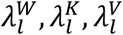 are inverse learning rate factors that express the insensitivity of synaptic strengths to plasticity. Please refer to [17] for details.

### Numerical simulations

Here, we demonstrate the performance of canonical neural networks using maze tasks—as an example of a delayed reward task. The agent comprised the aforementioned canonical neural networks (**Fig. 4a**). Thus, it implicitly performs active inference by minimising variational free energy. The maze affords a discrete state space (**Fig. 4b**). The agent received the states of the neighbouring cells as sensory inputs, and its neural activity represented the hidden states (**Fig. 4a**, panels on the right; See Methods E for further details). Although we denoted *s* as hidden states, the likelihood mapping *A* was a simple identity mapping in these simulations. When solving the above equations (6), (8), and (9), the agent’s neural network implicitly updates posterior beliefs about its behaviour based on the policy mapping. It then selects an appropriate action to move towards a neighbouring cell according to the inferred policy. The action was accepted if the selected movement was allowed.

Before training, the agent moved to a random direction in each step, resulting in a failure to reach the goal position (right end) within the time limit. During training, the neural network updated synaptic strengths depending on its neural activity and ensuing outcomes (i.e., risk). The training comprised a cycle of action and learning phases. In the action phase, the agent enacted a sequence of decisions, until it reached the goal or *T* = 2 × 10^4^ time steps passed (**Fig. 4c**). In the learning phase, the agent evaluated the risk associated with past decisions after a certain period: the risk was minimum (i.e., Γ(*t*) = 0) if the agent moved rightwards with a certain distance during the period; otherwise Γ(*t*) = 0.45 if the agent moved rightwards during the period, or Γ(*t*) = 0.55 if it did not. The synaptic strengths *V* (i.e., the policy mapping) were then potentiated if the risk was low, or suppressed otherwise, based on equation (9). This mechanism made it possible to optimise decision making. Other synapses (*W*, *K*) were also updated based on equation (8), although we assumed a small learning rate to focus on the implicit policy learning. Through training, the neural network of the agent self-organised its behaviour to efficiently secure its goal (**Fig. 4d**).

With this set up in place, we numerically validated the dependency of performance on the threshold factors (*ϕ*, φ). Consistent with our theoretical prediction—that *ϕ* and φ encode prior beliefs about hidden states (*D*) and decisions (*E*)—alternations of φ = ln *E* from the optimum to a suboptimal value changed the landscape of the cost function (i.e., variational free energy), thereby providing suboptimal inferences and decisions (in relation to the environment). Subsequently, the suboptimal network firing thresholds led to a suboptimal behavioural strategy, taking a longer time or failing to reach the goal (**Fig. 4e**). Thus, we could attribute the agent’s impaired performance to its suboptimal priors. This treatment renders neural activity and adaptive behaviours of the agent highly explainable and manipulatable in terms of the appropriate prior beliefs—implicit in firing thresholds—for a given task or environment. In other words, these results suggest that firing thresholds are the neuronal substrates that encode state and decision priors, as predicted mathematically.

**Figure 4.**
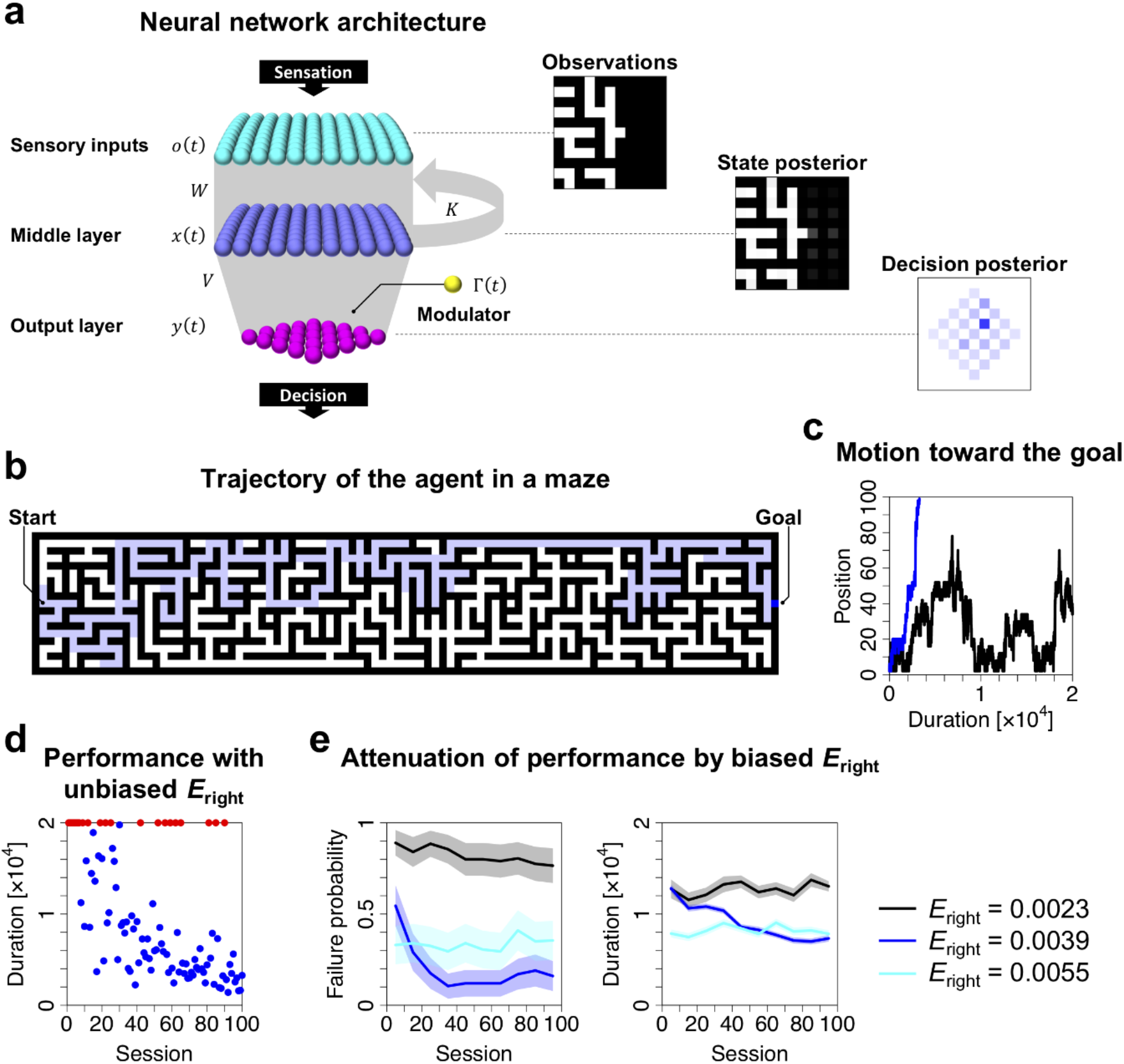
Simulations of neural networks solving maze tasks. (**a**) Neural network architecture. The agent receives the states (pathway or wall) of the neighbouring 11 × 11 cells as sensory inputs. A policy decision represents a four-step sequence of actions (selected from up, down, left, or right), resulting in 256 options in total. The panels on the right depict observations and posterior beliefs about hidden states and decisions. (**b**) General view of the maze. The maze comprises a discrete state space, wherein white and black cells indicate pathways and walls, respectively. A thick blue cell indicates the current position of the agent, while the thin blue line is its trajectory. Starting from the left, the agent needs to reach the right edge of the maze within *T* = 2 × 10^4^ time steps. (**c**) Trajectories of the agent’s x-axis position in sessions before (black, session 1) and after (blue, session 100) training. (**d**) Duration to reach the goal when the neural network operates under uniform decision priors *E*_right_ = *E*_left_ = *E*_up_ = *E*_down_ = 1/256 ≈ 0.0039 (where *E*_right_ indicates the prior probability to select a decision involving the rightward motion in the next step). Blue and red circles indicate succeeded and failed sessions, respectively. (**e**) Failure probability (left) and duration to reach the goal (right) when the neural network operates under three different prior conditions *E*_right_ = 0.0023,0.0039,0.0055, where *E*_left_ = 0.0078 − *E*_right_ and *E*_up_ = *E*_down_ = 0.0039 hold. The line indicates the average of 10 successive sessions. Although the neural network with *E*_right_ = 0.0055 exhibits better performance in the early stage, it turns out to overestimate a preference of the rightward motion in later stages, even when it approaches the wall. (e) was obtained with 20 distinct, randomly generated mazes. Shaded areas indicate the standard error. Refer to Methods E for further details.

Furthermore, when the updating of *ϕ* and φ is slow in relation to experimental observations, *ϕ* and φ can be estimated through Bayesian inference based on empirically observed neuronal responses (see Methods E for details). Using this approach, we estimated implicit prior *E*—which is encoded by φ—from sequences of neural activity generated from the synthetic neural networks used in the simulations reported in **Fig. 4**. We confirmed that the estimator was a good approximation to the true *E* (**Fig. 5a**). The estimation of *ϕ* and φ based on empirical observations offered the reconstruction of the cost function (i.e., variational free energy) that an agent employs. The resulting cost function could predict subsequent learning of behaviours within previously unexperienced, randomly generated mazes—without observing neural activity and subsequent behaviour (**Fig. 5b**). This is because—given the canonical neural network at hand—the learning self-organisation is based exclusively on state and decision priors, implicit in *ϕ* and φ. Therefore, the identification of these implicit priors is sufficient to asymptotically determine the fixed point of synaptic strengths when *t* is large (see Methods C, D for further details; see also [17]). These results highlight the utility of the proposed equivalence to understand neuronal mechanisms underlying adaptation of neural activity and behaviour through accumulation of past experiences and ensuing outcomes.

**Figure 5.**
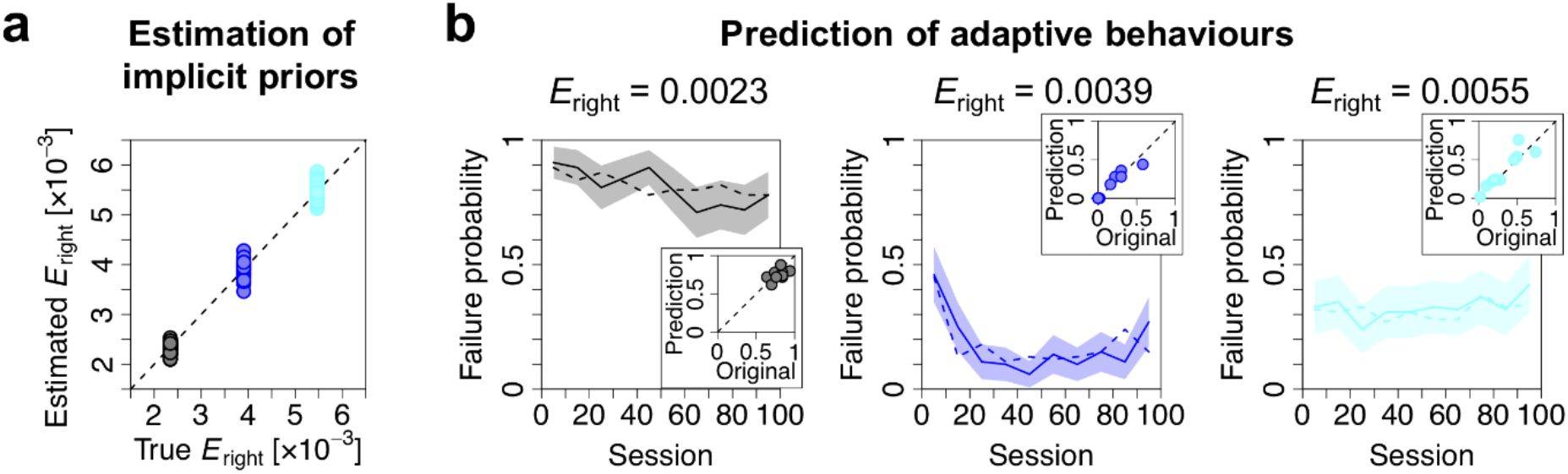
Estimation of implicit priors enables the prediction of subsequent learning. (**a**) Estimation of implicit prior *E*_right_—encoded by threshold factor φ—under three different prior conditions (black, blue, and cyan; c.f., **Fig. 4**). Here, φ was estimated through Bayesian inference based on sequences of neural activity, obtained with 10 distinct mazes. Then, *E*_right_ was computed by ln *E* = φ for each of 64 elements. The other 192 elements of *E* (i.e., *E*_left_, *E*_up_, *E*_down_) were also estimated. (**b**) Prediction of the learning process within previously unexperienced, randomly generated mazes. Using the estimated *E*, we reconstructed the computational architecture (i.e., neural network) of the agent. Then, we simulated the adaptation process of the agent’s behaviour using the reconstructed neural network, and computed the trajectory of the probability of failure to reach the goal within *T* = 2 × 10^4^ time steps. The resulting learning trajectories (solid lines) predict the learning trajectories of the original agent (dashed lines) under three different prior conditions, in the absence of observed neural responses and behaviours. Lines and shaded areas indicate the mean and standard error, respectively. Inset panels depict comparisons between the failure probability of the original and reconstructed agent after learning (average over session 51–100), within 10 previously unexperienced mazes. Refer to Methods E for further details.

## DISCUSSION

Biological organisms formulate plans to minimise future risk. In this work, we captured this characteristic in biologically plausible terms under minimal assumptions. We derived simple differential equations that can be plausibly interpreted in terms of a neural network architecture that entails degrees of freedom with respect to certain free parameters (e.g., firing threshold). These free parameters play the role of prior beliefs in variational Bayesian formation. Thus, the accuracies of inferences and decisions depend upon prior beliefs, implicit in neural networks. Consequently, synaptic plasticity with false prior beliefs lead to suboptimal inferences and decisions for any task under consideration.

A simple Hebbian plasticity strengthens synaptic wiring when pre- and post-synaptic neurons fire together, which enhances the association between (pre-synaptic) causes and (post-synaptic) consequences [23]. Hebbian plasticity depends on the activity level [24,25], spike timings [31,32], or burst timings [33] of pre- and post-synaptic neurons. Furthermore, modulatory factors can regulate the magnitude and parity of Hebbian plasticity, possibly with some delay in time, leading to the emergence of various associative functions [26–28]. These modulations have been observed empirically with various neuromodulators and neurotransmitters, such as dopamine [30,34,35], noradrenaline [36,37], muscarine [38], and GABA [39,40], as well as glial factors [41].

In particular, a delayed modulation of synaptic plasticity is well-known with dopamine neurons [30]. We mathematically demonstrated that such a plasticity enhances the association between the pre-post mapping and the future value of the modulatory factor, where the latter is cast as a risk function. This means that post-synaptic neurons self-organise to react in a manner that minimises future risk. Crucially, this computation corresponds formally to variational Bayesian inference under a particular form of POMDP generative models, suggesting that the delayed modulation of Hebbian plasticity is a realisation of active inference. Regionally specific projections of neuromodulators may allow each brain region to optimise activity to minimise risk, and leverage a hierarchical generative model implicit in cortical and subcortical hierarchies. This is reminiscent of theories of neuromodulation and (meta-)learning developed previously [42]. Our work may be potentially useful, when casting these theories in terms of generative models and variational free energy minimisation.

It is remarkable that the proposed equivalence can be leveraged to identify a generative model that an arbitrary neural network implicitly employs. This contrasts with naïve neural network models that address only the dynamics of neural activity and plasticity. If the generative model differs from the true generative process—that generates the sensory input—inferences and decisions are biased (i.e., suboptimal), relative to Bayes optimal inferences and decisions based on the right sort of prior beliefs. In general, the implicit priors may or may not be equal to the true priors; thus, a generic neural network is typically suboptimal. Nevertheless, these implicit priors can be optimised by updating free parameters (e.g., threshold factors *ϕ*, φ) based on the gradient descent on cost function *L*. By updating the free parameters, the network will eventually, in principle, becomes Bayes optimal for any given task. In essence, when the cost function is minimised with respect to neural activity, synaptic strengths, and any other constants that characterise the cost function, the cost function becomes equivalent to variational free energy with the optimal prior beliefs. Simultaneously, the expected risk is minimised because variational free energy is minimised only when the precision of the risk (*γ*_*t*_) is maximised (see Methods A for further details).

When the rate coding activation function differs from the sigmoid function, it can be assumed that neurons encode state posteriors under a generative model that differs from a typical POMDP model considered in this work. Nevertheless, the complete class theorem guarantees the existence of some pair of generative model (i.e., priors) and cost function that correspond to an arbitrary activation function. The form or time-window of empirically observed plasticity rules can also be used to identify the implicit cost and risk functions—and further to reverse engineer the task or problem that the neural network is solving or learning: c.f., inverse reinforcement learning [43]. In short, neural activity and plasticity can be interpreted, universally, in terms of Bayesian belief updating.

The class of neural networks we consider can be viewed as a class of reservoir networks [44,45]. The proposed equivalence could render such reservoir networks explainable—and may provide the optimal plasticity rules for these networks to minimise future risk—by using the formal analogy to variational free energy minimisation (under the particular form of POMDP models). A clear interpretation of reservoir networks remains an important open issue in computational neuroscience and machine learning.

The equivalence between neural network dynamics and gradient flows on variational free energy is empirically testable using electrophysiological recordings or functional imaging of brain activity. We have previously shown that the self-organisation of *in vitro* neural networks minimises empirically computed variational free energy in a manner consistent with variational free energy minimisation under a POMDP generative model [46,47]. Our analyses in the present work speak to the predictive validity of the proposed formulation: when the threshold factors (*ϕ*, φ) can be treated as constants—during a short experiment—we obtain the analytical form of fixed points for synaptic update rules (Methods D). Furthermore, *ϕ* and φ can be estimated using empirical data (Methods E). This approach enables the reconstruction of the cost function and prediction of subsequent learning process, as demonstrated in **Fig. 5** using *in silico* data. Hence, it is possible to examine the predictive validity of the proposed theory by comparing the predicted synaptic trajectory with the actual trajectory. In future work, we hope to address these issues using *in vitro* and *in vivo* data.

In summary, a class of cost functions for canonical neural networks can be cast as variational free energy. Formal correspondences exist between priors, posteriors, and cost functions. This means that canonical neural networks that optimise their cost functions implicitly perform active inference. This approach enables identification of the implicit generative model and reconstruction of variational free energy that neural networks employ. This means, in principle, neural activity, behaviour, and learning through plasticity can be predicted under Bayes optimality assumptions.

## Data Availability

All relevant data are within the paper. The MATLAB scripts are available at https://github.com/takuyaisomura/reverse_engineering.

## Acknowledgements

This work was supported in part by the grant of Joint Research by the National Institutes of Natural Sciences (NINS Program No. 01112005). T.I. is funded by RIKEN Center for Brain Science. H.S. is funded by MEXT/JSPS KAKENHI Grant Number JP 20K11709. K.J.F. is funded by a Wellcome Principal Research Fellowship (Ref: 088130/Z/09/Z). The funders had no role in study design, data collection and analysis, decision to publish, or preparation of the manuscript.

## METHODS

### A. Generative model

The proposed POMDP model comprises *N*_*S*_-dimensional hidden states 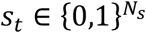 that depend on the previous states *S*_*t*-1_ through a transition probability of *B*^δ^, and a process of generating *N*_*O*_-dimensional observations 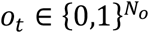 from those states through a likelihood mapping *A* (**Fig. 2**). Here, the transition probability *B*^δ^ is a function of *N*_f_-dimensional decisions of an agent *δ*_*t*_ ∈ {0,1}*N*^δ^, indicating that the agent’s behaviour changes the subsequent states of the external milieu. Each state, observation, and decision take the values 1 or 0. We use *o*_1:*t*_ = {*o*_1_,…, *o*_*t*_} to denote a sequence of observations. Hereafter, *i* indicates the *i*-th observation, *j* indicates the *j*-th hidden state, and *k* indicates the *k*-th decision.

Due to the multi-dimensional (i.e., factorial) nature of the states, *A* and *B* (and *C*) are usually the outer products of sub matrices (i.e., tensors); see also [17]. The probability of an observation is determined by the likelihood mapping, from *S*_*t*_ to 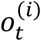, in terms of a categorical distribution: 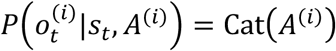, where the elements of *A*^(*i*)^ are given by 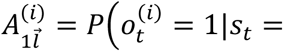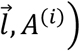 and 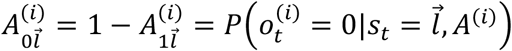. This encodes the probability of 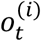 takes 1 or 0 when 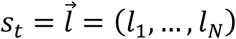. The prior belief of 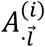 is defined by Dirichlet distribution 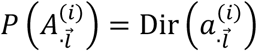 with concentration parameter 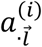. The hidden states are determined by the transition probability, from *S*_*t*-1_ to *S*_*t*_, depending on a given decision, in terms of a categorical distribution: 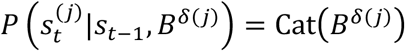, where the elements of *B*^δ(*j*)^ are given by 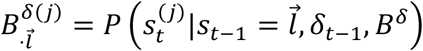. The prior distribution of 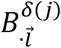 is defined by Dirichlet distribution 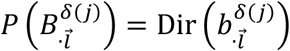 with concentration parameter 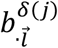.

The policy mapping is optimised based on a generative model of decisions *δ*_*t*_ conditioned on the current risk *γ*_*t*_ ∈ {0,1} that obeys 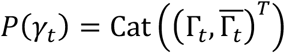, where 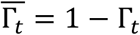. This can be viewed as a postdiction of past decisions. We express the conditional probability of 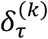 as a form of a mixture model with respect to 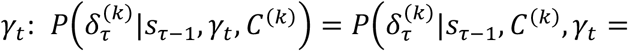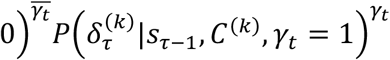. Namely, the agent assumes (i.e., postdicts) that 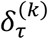 was sampled by 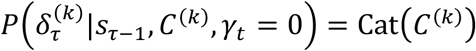 after receiving *γ*_*t*_ = 0, or sampled by 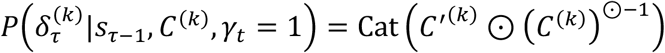 after receiving *γ*_*t*_ = 1. Because the agent does not yet observe the consequence of current decision *δ*_*t*_, *δ*_*t*_ is sampled from 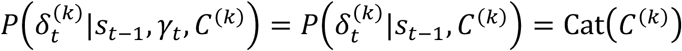 to minimise the future risk. The prior belief of 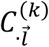 is defined by Dirichlet distribution 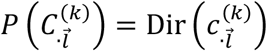 with concentration parameter 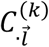.

Therefore, when observing *γ*_*t*_ = 0, the agent regards that the past decisions were sampled from the preferable policy mapping *C* that minimises Γ_*t*_, and thereby updates the posterior belief of *C* to facilitate the association between *δ*_2_ and *S*_τ-1_. In contrast, when observing *γ*_*t*_ = 1, the agent hypothesises that this is because the past decisions were sampled from the unpreferable policy mapping, and thereby updates the posterior belief of *C* to reduce (forget) the association. In short, this postdiction evaluates the past decisions after observing their consequence. The posterior belief of *C* is then updated by associating the past decision rule (policy) and current risk, leading to the optimisation of decisions to minimise future risk.

An advantage of this generative model—based on counterfactual causality—is that the agent does not need to explicitly compute the expected future risk based on the current states, because it instead updates the policy mapping *C*, by associating the current risk with past decisions. Note that this construction of risk corresponds to a simplification of expected free energy, that would normally include risk and ambiguity, where risk corresponds to the Kullback-Leibler divergence between the posterior predictive and prior distribution over outcomes [12,13]. However, by using a precise likelihood mapping, ambiguity can be discounted and expected free energy reduces to the sort of decision risk considered in this work.

We can now define the generative model as equation (2), where *P*(*S*_1_|*S*_0_, *δ*_0_, *B*) = *P*(*S*_1_) = Cat(*D*) and *P*(*δ*_1_|*S*_0_, *γ*_*t*_, *C*) = *P*(*δ*_1_) = Cat(*E*) are assumed. We further suppose that *δ*_τ_, given *S*_τ-1_, is conditionally independent of {*o*_τ_, *S*_τ_}, and that only the generation of *δ*_τ_ depends on *γ*_*t*_, as visualised in the factor graph (**Fig. 2**). Equation (2) can be further expanded as follows:

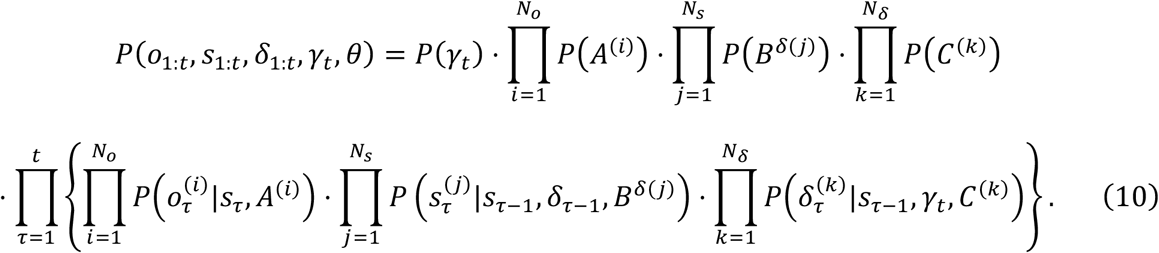

As described in the Results section, this form of generative model is suitable to characterise a class of canonical neural networks defined by equation (6). This means that none of aforementioned assumptions about the generative model limits the scope of the proposed equivalence between neural networks and variational Bayesian inference, as long as neural networks satisfy assumptions 1–3.

### B. Variational free energy

The agent aims to minimise surprise, or equivalently maximise the marginal likelihood of outcomes, by minimising variational free energy as a tractable proxy. Thereby, they perform approximate or variational Bayesian inference. From the above-defined generative model, we motivate a mean-field approximation to the posterior distribution as follows:

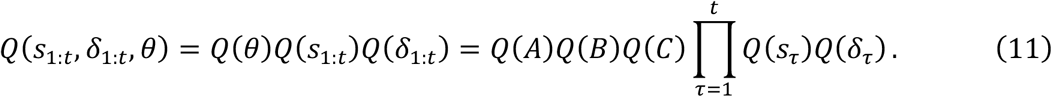

Here, the posterior beliefs of *S*_τ_ and *δ*_τ_ are categorical distributions, *Q*(*S*_τ_) = Cat(**s**_τ_) and *Q*(*δ*_τ_) = Cat(**δ**_τ_), respectively. Whereas, the posterior beliefs of *A*, *B*, and *C* are Dirichlet distributions, *Q*(*A*) = Dir(**a**), *Q*(*B*) = Dir(**b**), and *Q*(*C*) = Dir(**c**), respectively. In this expression, **s**_τ_ and **δ**_τ_ represent the expectations between 0 and 1, and **a**, **b**, and **c** express the (positive) concentration parameters.

In this paper, the posterior transition mapping is averaged over all possible decisions, **B** = E_*Q*(*δ*)_[**B**^δ^], to ensure exact correspondence to canonical neural networks. Moreover, we suppose that **A** comprises the outer product of sub-matrices **A**^(*i,j*)^ ∈ ℝ^2×2^ to simplify calculation of the posterior beliefs, i.e., 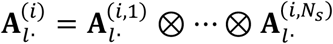 for *l* = 0,1. We also suppose that **B** and **C** comprise the outer products of sub-matrices 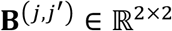 and **C**^(*k,j*)^ ∈ ℝ^2×2^, respectively. The expectation over the parameter posterior 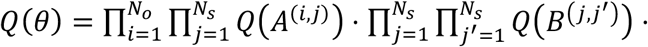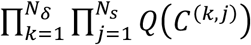 is denoted as E_*Q(*θ)_[∙] ≔ ∫∙ *Q*(*θ*)*dθ*. Using this, the posterior expectation of a parameter 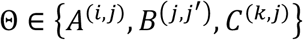 is expressed using the corresponding concentration parameter 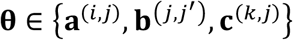 as follows:

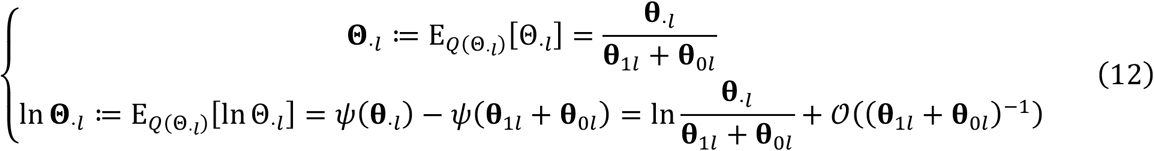

for *l* = 0,1, where φ(∙) is the digamma function.

In terms of decisions, because *P*(*δ*_τ_|*S*_τ-1_, *C*, *γ*_*t*_ = 1) ∝ Cat(*C*^⊙-1^) in this setup, the complexity associated with past decision is given by 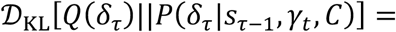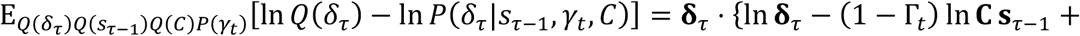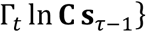 for 1 ≤ *τ* ≤ *t* − 1, up to the *C*′-dependent term which is negligible when computing the posterior beliefs. Whereas, the current decision is made to minimise the complexity 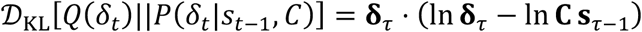.

Variational free energy is defined as a functional of the posterior beliefs, given as:

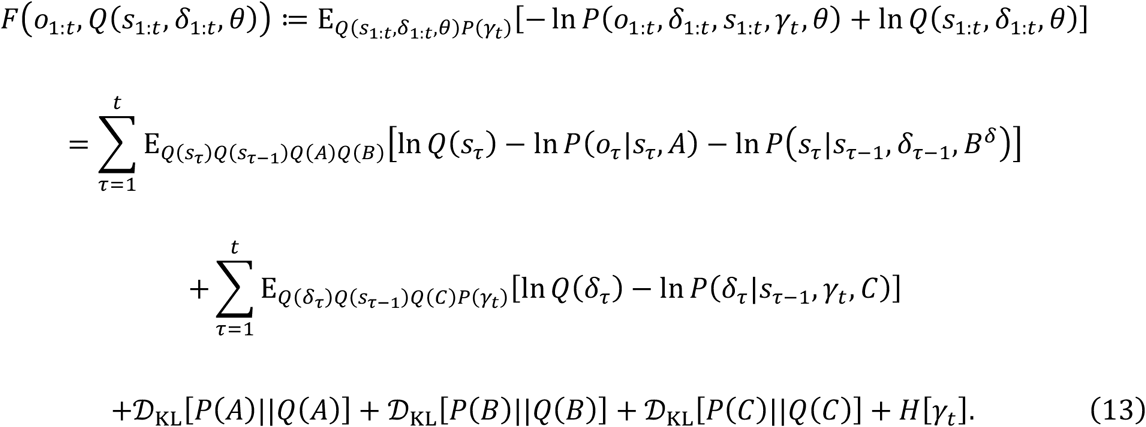

This provides an upper bound of sensory surprise − ln *P*(*o*_1:*t*_). Here, 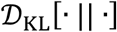 is complexity of parameters scored by the Kullback-Leibler divergence. Minimisation of variational free energy is attained when the entropy of the risk, 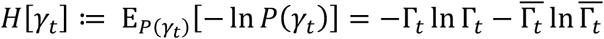, is minimised. This is achieved when Γ_*t*_ shifts toward 0, meaning that the risk minimisation is a corollary of variational free energy minimisation (the case where Γ_*t*_ shifts toward 1 is negligible). Under the MDP scheme, *F* is expressed as a function of the posterior expectations, *F* = *F*(*o*_1:*t*_, **s**_1:*t*_, **δ**_1:*t*_, **θ**). Thus, using the vector expression, variational free energy under our POMDP model is provided as follows:

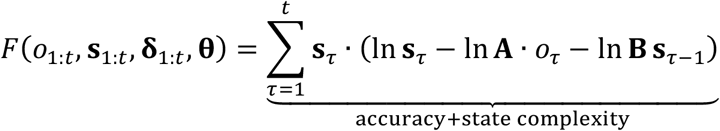

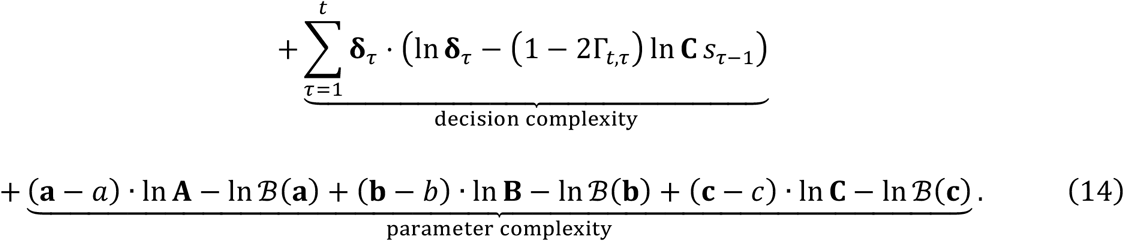

Here, 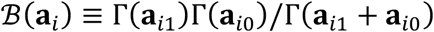 is the beta function. Here, ln **A** ∙ *o*_τ_ indicates the inner product of ln **A** and one-hot expressed *o*_τ_, which is a custom to express the sum of the product of 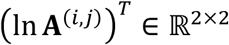 and 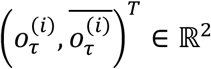 over all *i*.

The first and second terms of equation (14)—comprising accuracy and the complexity of state and decision—increases in proportion to time *t*. Conversely, other terms—the complexity of parameters—increases in the order of ln *t*, which is thus negligible when *t* is large. Thus, we will drop the parameter complexity by assuming that the scheme has experienced a sufficient number of observations. Please see [17] for further details. The entropy of the risk is omitted as it is of order 1.

Based on the Bayes theorem, *P*(*S*_τ_|*S*_τ-1_, *B*^δ^) ∝ *P*(*S*_τ-1_|*S*_τ_, *B*^δ^)*P*(*S*_τ_) and *P*(*δ*_τ_|*S*_τ-1_, *γ*_*t*_, *C*) ∝ *P*(*S*_τ-1_|*δ*_τ_, *γ*_*t*_, *θ*)*P*(*δ*_τ_) hold, where *P*(*S*_τ-1_) is supposed to be a flat prior belief. Thus, the inverse transition and policy mappings are given as **B**^✝^ = **B**^*T*^diag[*D*]^-1^ and **C**^✝^ = **C**^*T*^diag[*E*]^-1^, respectively. Thus, **s**_τ_ ⋅ ln **B s**_τ-1_ = **s**_τ_ ⋅ (ln **B**^✝^ ⋅ **s**_τ-1_ + ln *D*) and **δ**_τ_ ⋅ ln **C s**_τ-1_ = **δ**_τ_ ⋅ (ln **C**^✝^ ⋅ **s**_τ-1_ + ln *E*) hold. Accordingly, equation (14) becomes

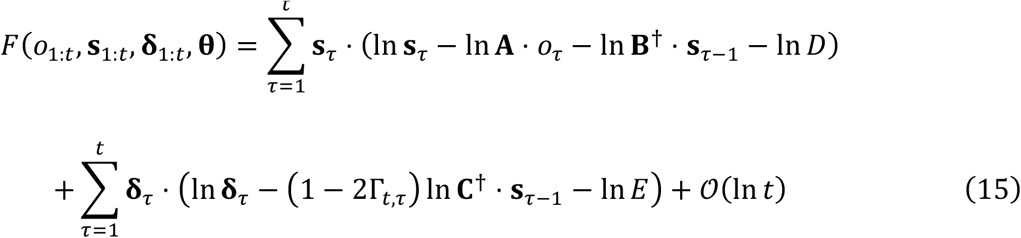

as expressed in **Fig. 3**(top). Here, the prior beliefs about states and decisions, *P*(*S*_τ_) = Cat(*D*) and *P*(*δ*_τ_) = Cat(*E*), alter the landscape of variational free energy. We will see below that this specific form of variational free energy corresponds formally to a class of cost functions for canonical neural networks.

### C. Inference and learning

Inference optimises the posterior beliefs about the hidden states and decisions by minimising variational free energy. The posterior beliefs are updated by the gradient descent on 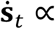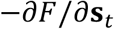 and 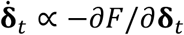. The fixed point of these updates furnishes the posterior beliefs, which are analytically computed by solving ∂*F*/∂**s**_*t*_ = 0 and ∂*F*/∂**δ**_*t*_ = 0. Thus, from equation (15), the posterior belief about the hidden states is provided as follows:

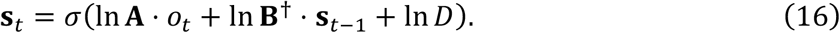

Moreover, the posterior belief about the decisions is provided as follows:

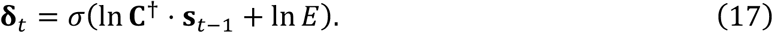

Here, σ(∙) denotes the softmax function; and *D* and *E* denote the prior beliefs about hidden states and decisions, respectively, which we assume are fixed in this paper. Note that equations (16) and (17) are equivalent to **s**_*t*_ = σ(ln **A** ⋅ *o*_*t*_ + ln **B s**_*t*-1_) and **δ**_*t*_ = σ(ln **C s**_*t*-1_), respectively, as **B**^✝^ = **B**^*T*^diag[*D*]^-1^ and **C**^✝^ = **C**^*T*^diag[*E*]^-1^. Notably, 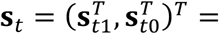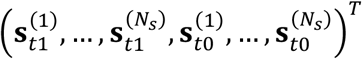 indicates a block column vector of the state posterior under a mean-field assumption, where 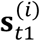 is the posterior belief that 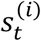 takes a value of one. Since 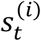 takes either one or zero, a binary value, 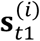 has the form of a sigmoid function. Here, we assume that only the state posterior **s**_*t*_ at the latest time is updated at each time *t*; thus, no message pass exists from **s**_*t*+1_ to **s**_*t*_. The state posterior at time *t*–1 is retained in the previous value. This treatment corresponds to the Bayesian filter, as opposed to the smoother usually considered in active inference schemes.

Equations (16) and (17) are analogue to a two-layer neural network that entails recurrent connections in the middle layer. In this analogy, **s**_*t*1_ and **δ**_*t*1_ are viewed as the middle- and output-layer neural activity, respectively. Moreover, ln **A** ⋅ *o*_*t*_, ln **B**^✝^ ⋅ **s**_*t*-1_, and ln **C**^✝^ ⋅ **s**_*t*-1_ corresponds to synaptic inputs, and ln *D* and ln *E* relates to firing thresholds. These priors and posteriors turn out to be identical to the components of canonical neural networks, as described in the Results and Methods D.

Furthermore, learning optimises the posterior beliefs about the parameters **θ** = {**a**, **b**, **c**} by minimising variational free energy. The posterior beliefs are updated by the gradient descent on *F*, 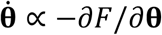. By solving the fixed point ∂*F*/∂**θ** = *O* of equation (14), the posterior beliefs about parameters are provided as follows:

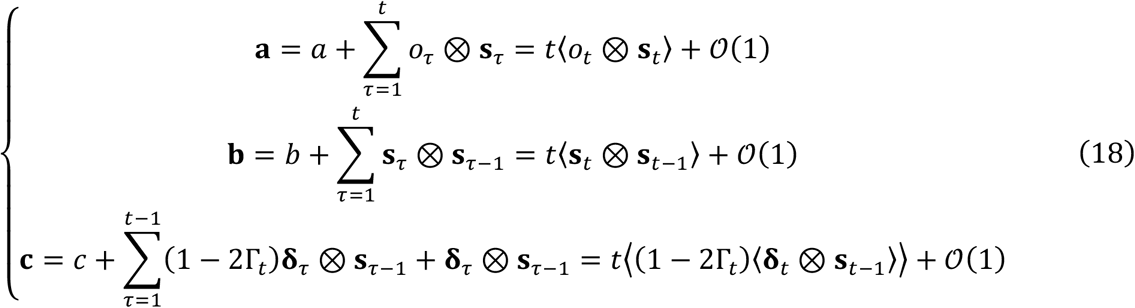

Note that ⨂ denotes the outer product operator, and 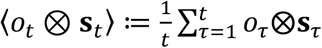. Here, *a*, *b*, *c* are the prior beliefs, which are of order 1 and thus negligibly small relative to the leading order term when *t* is large. Thus, the posterior expectations of any parameters Θ (i.e., **Θ** and ln **Θ**) are obtained using equations (12) and (18). These parameter posteriors turn out to correspond formally to synaptic strengths (*W*, *K*, *V*) owing to the equivalence of variational free energy and neural network cost function.

### D. Neural networks

Updates of neural activity are defined by equation (6). When the time constant of neural activity is smaller than that of sensory inputs, the fixed point of equation (6)—i.e., *x* and *y* that give 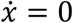 and 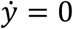—is provided as follows:

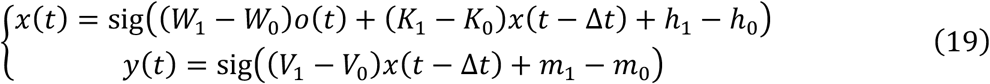

The adaptive firing thresholds are given as functions of synaptic strengths, 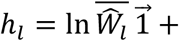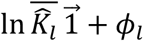 and 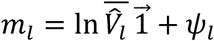 (for *l* = 0,1), where exp(*ϕ*_1_) + exp(*ϕ*_0_) = 1 and exp(φ_1_) + exp(φ_0_) = 1 hold. The form of equation (19) is identical to the state and decision posteriors in equations (16) and (17) of the variational Bayesian formation.

Considering that neural activity corresponds to the posterior beliefs about states and decisions, one might consider the relationship between synaptic strengths and parameter posteriors. As we mathematically demonstrated in the Results section, owing to the equivalence between variational free energy *F* and the neural network cost function *L*, i.e., equation (5) versus equation (7), synaptic strengths correspond formally to parameter posteriors. The ensuing synaptic update rules—derived as the gradient descent on *L*—are expressed in equations (8) and (9). They have a biologically plausible form, comprising Hebbian plasticity accompanied with an activity-dependent homeostatic term. The product of Γ(*t*) and the associated term modulates plasticity depending on the quality of past decisions, after observing their consequences, leading to minimisation of the future risk. In other words, the modulation by Γ(*t*) represents a postdiction that the agent implicitly conducts, wherein the agent regards its past decisions as preferable when Γ(*t*) is low and memorises the strategy. Conversely, it regards them as unpreferable when Γ(*t*) is high and forgets the strategy.

In particular, when *ϕ* and φ are constants, the fixed point of synaptic strengths that minimise *L* is expressed analytically as follows: for simplification, we employ the notation using ω_*i*_, *pre*_*i*_, *post*_*i*_, and *n*_*i*_, as described in the Results section. The derivative of firing threshold *n*_*i*_ with respect to synaptic strength matrix ω_*i*_ yields the sigmoid function of ω_*i*_, i.e., 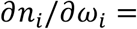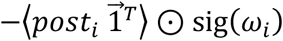. Here, ⊙ indicates the element-wise product (Hadamard product). The fixed point of synaptic strengths ensures ∂*L*/∂ω_*i*_ = *O*. Thus, from equations (8) and (9), it is analytically expressed as

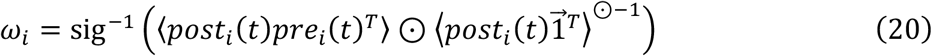

for *i* = 1,…,4, and

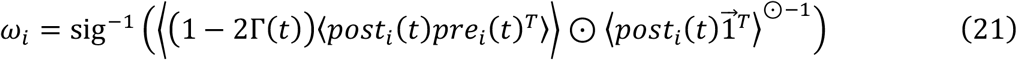

for *i* = 5,6. Equations (20) and (21) correspond to the posterior belief about parameters *A*, *B*, *C*, which are shown in equation (18).

### E. Protocols for numerical simulations and data analyses

In **Fig. 4**, the neural network of the agent was characterised by a set of internal states *φ* = {*x*, *y*, *W*, *K*, *V*, *ϕ*, φ}. Neural activity (*x*, *y*) was updated by following equation (6), while synaptic strengths (*W*, *K*, *V*) were updated by following equations (8) and (9). Here, we supposed that neural activity converges quickly to the steady state relative to the change of observations. This treatment allowed us to compute the network dynamics based on equations (19)–(21), which could reduce the computational cost for numerical simulations. Synaptic strengths *W* were initialised as a matrix sufficiently close to the identity matrix; whereas, synaptic strengths *K* and *V* were initialised as matrices with uniform values. This treatment served to focus on the policy learning implicit in the update of *V*. The threshold factors (*ϕ*, φ), which encoded the prior beliefs about hidden states (*D*) and decisions (*E*), were pre-defined and fixed over the sessions. In **Fig. 4e**, we varied *E* to show how performance depends on these priors. Namely, *E*_1_ = (*E*_right_,…, *E*_right_,*E*_left_,…, *E*_left_, *E*_up_,…, *E*_up_,*E*_down_,…, *E*_down_)^*T*^ ∈ [0,1]^256^ was characterised by four values *E*_right_, *E*_left_, *E*_up_, *E*_down_ ∈ [0,1], where *E*_right_ indicates the prior probability to select a decision involving the rightward motion in the next step.

When the belief updating of implicit priors (*D*, *E*) is slow in relation to experimental observations, *D* and *E*—which are encoded by *ϕ* and φ—can be viewed as being fixed over a short period of time, as an analogy to a homeostatic plasticity over longer time scales [51]. In this case, through variational free energy minimisation based on empirically observed neuronal responses, the estimators of *ϕ* and φ are obtained as follows:

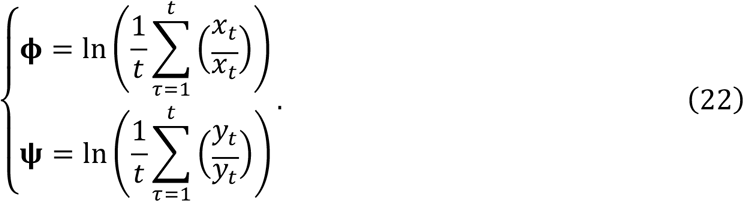

Note that we suppose the constraints 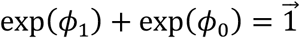 and 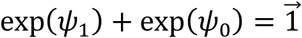. This characterisation finessed the estimation of implicit priors (**Fig. 5a**) and the reconstruction of variational free energy. Furthermore, the reconstructed variational free energy was used to predict subsequent inference and learning, without observing neural activity (**Fig. 5b**). In **Fig. 5**, for simplicity, the form of the risk function was supposed to be known when reconstructing the cost function. Although we did not estimate *ϕ* in **Fig. 5**, the previous work showed that our approach can estimate *ϕ* from simulated neural activity data [17].

## Footnotes

* Although, by convention, active inference uses *C* to denote the prior preference, this work uses *C* to denote a mapping to determine a decision depending on the previous state. Herein, the prior preference is implicit in the risk function Γ_*t*_. Due to construction, *C*′ does not explicitly appear in the inference; thus, it is omitted in the following formulations.

